# Photon yield enhancement of red fluorophores at cryogenic temperatures

**DOI:** 10.1101/263848

**Authors:** Christiaan N. Hulleman, Weixing Li, Ingo Gregor, Bernd Rieger, Jörg Enderlein

## Abstract

Single Molecule Localization Microscopy has become one of the most successful and widely applied methods of Super-resolution Fluorescence Microscopy. Its achievable resolution strongly depends on the number of detectable photons from a single molecule until photobleaching. By cooling a sample from room temperature down to liquid nitrogen temperatures, the photostability of dyes can be enhanced by more than 100 fold, which results in an improvement in localization precision greater than 10 times. Here, we investigate a variety of fluorescent dyes in the red spectral region, and we find an average photon yield between 3.5 · 10^6^ to 11 · 10^6^ photons before bleaching at liquid nitrogen temperatures, corresponding to a theoretical localization precision around 0.1 nm.

## 1 Introduction

The last 25 years have seen a revolution in optical far field microscopy, in particular due to the invention of super-resolution methods such as STED, PALM and STORM [1]. Especially the two latter methods, which are methods of the class of Single-Molecule Localization Microscopy (SMLM), now routinely achieve image resolutions of 10-20 nm [2, 3]. The attainable localization precision of SMLM is ultimately limited by the amount of detected photons until bleaching (or photoswitching), and follows the generic law

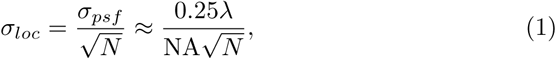

where λ is the emission wavelength, NA the numerical aperture of the objective, and *N* is the number of detected photons [4]. More complex equations have been developed taking background and camera pixelation into account [5], but equation 1 represents the most favorable scenario and thus a lower limit for the attainable resolution.

As can be seen, one way to increase the achievable localization accuracy and thus image resolution is to increase the NA. However, another way is to increase the number of detected photons *N*. Thus, by cooling a sample and thereby increasing the photostability of the fluorescent dye molecules can have a tremendous impact on resolution in SMLM. Cooling reduces all photochemical reaction rates and thus inhibits bleaching, thereby enabling significantly more photons to be detected from a molecule, resulting in a localization precision in the Ångström range [6, 7]. Weisenburger et al. compared the performance of dyes with an emission peak of 552-576 nm at liquid helium temperatures (4.4 K), finding an increase of one to two orders of magnitude in the number of emitted photons [6]. Li et al. built a cryo-fluorescence microscope with samples contained in a liquid nitrogen cryostat and analyzed the performance of ATTO 647N, finding an increase of more than two orders of magnitude in photon yield [7]. One more advantage of cryo-fluorescence microscopy is that it offers the possibility for correlative light and electron microscopy (cryo-CLEM), adding the specificity of fluorescent labeling to the superior structural resolution of electron microscopy (cryo-EM) [8, 9].

The resolution in a super-resolution image is not solely determined by the localization accuracy but equally by the density of emitters [10]. In cases where one can image many instances of a biological structure, the synthesis of many of these images into one final super-super-resolution image (particle averaging) can tremendously increase the effective label density and signal-to-noise ratio. In this case, the achievable resolution is finally only determined by the single-molecule localization precision *σ*_*loc*_ [11, 12]. Here, we see the biggest possible impact that improved localization precision at cryogenic temperatures can have on resolving molecular details with Ångström resolution.

For imaging of biological samples, red fluorescent dyes are particularly interesting due to the reduced auto-fluorescence and lower scattering intensity in the red spectral region [13]. Wurm et al. investigated the performance of red fluorophores at room temperature, comparing ATTO 647N and various Abberior STAR dyes [14]. ATTO 647N and Abberior STAR635 yielded the highest brightness, though the phosphorylated dye Abberior STAR635P had the best signal-to-noise ratio (SNR) due to reduced unspecific background staining. The high brightness of ATTO 647N and contrast of Abberior STAR635P makes them also excellent candidates for cryo-fluorescence microscopy.

In the present paper, we present experimental results for the photon yield of various red dyes at liquid nitrogen temperatures. In particular, we tested the dyes Abberior STAR635P, ATTO 647N, Alexa Fluor 647, ATTO 655, Cy5, and two Silicon-Rhodamine dyes, SiR and (Janelia Farm) JF 646. The intention is to find red dyes that offer the highest localization precision due to their increased photon yield at cryogenic temperatures.

## 2 Results and Discussion

For estimating the photostability of a dye, we immobilized a sparse concentration of the dye on a quartz surface and determined how many molecules could be localized in an image over the course of time. This number exponentially decayed with increasing time due to photobleaching. Thus, at room temperature we determined values of the half life, the time required to reduce the fluorescing molecules by 50%, ranging from *t*_1/2_ = 36 seconds for Abberior STAR635P to *t*_1/2_ = 173 seconds for ATTO 655 (Figure 1a). The excitation intensity for the room temperature measurement was 240 W/cm^2^. At cryogenic temperatures of ~89 K, each dye was measured at 3 different intensities, 100 W/cm^2^, 240 W/cm^2^, and 360 W/cm^2^ (Figure 1b-h). The half life was 128 times longer at cryogenic temperatures for ATTO 647N at an excitation intensity of 240 W/cm^2^ (*t*_1/2_ = 111 minutes compared to *t*_1/2_ = 52 seconds). The other dyes have a half life at cryogenic temperatures of *t*_1/2_ = 310 minutes for JF 646, *t*_1/2_ = 44 minutes for Alexa Fluor 647, *t*_1/2_ = 43 minutes for Cy5, *t*_1/2_ = 18 minutes for SiR, *t*_1/2_ = 12 minutes for Abberior STAR635P, and *t*_1/2_ = 10 minutes for ATTO 655, at the same excitation intensity of 240 W/cm^2^. Higher excitation intensities lead to faster bleaching of the fluorescent molecules; reducing the intensity to 100 W/cm^2^ decreases the bleaching rate whilst retaining a good signal-to-noise ratio in the images. The slow bleaching of JF 646, ATTO 647N, and Abberior STAR635P implies that they would be best for long-lasting cryogenic experiments.

**Figure 1.**
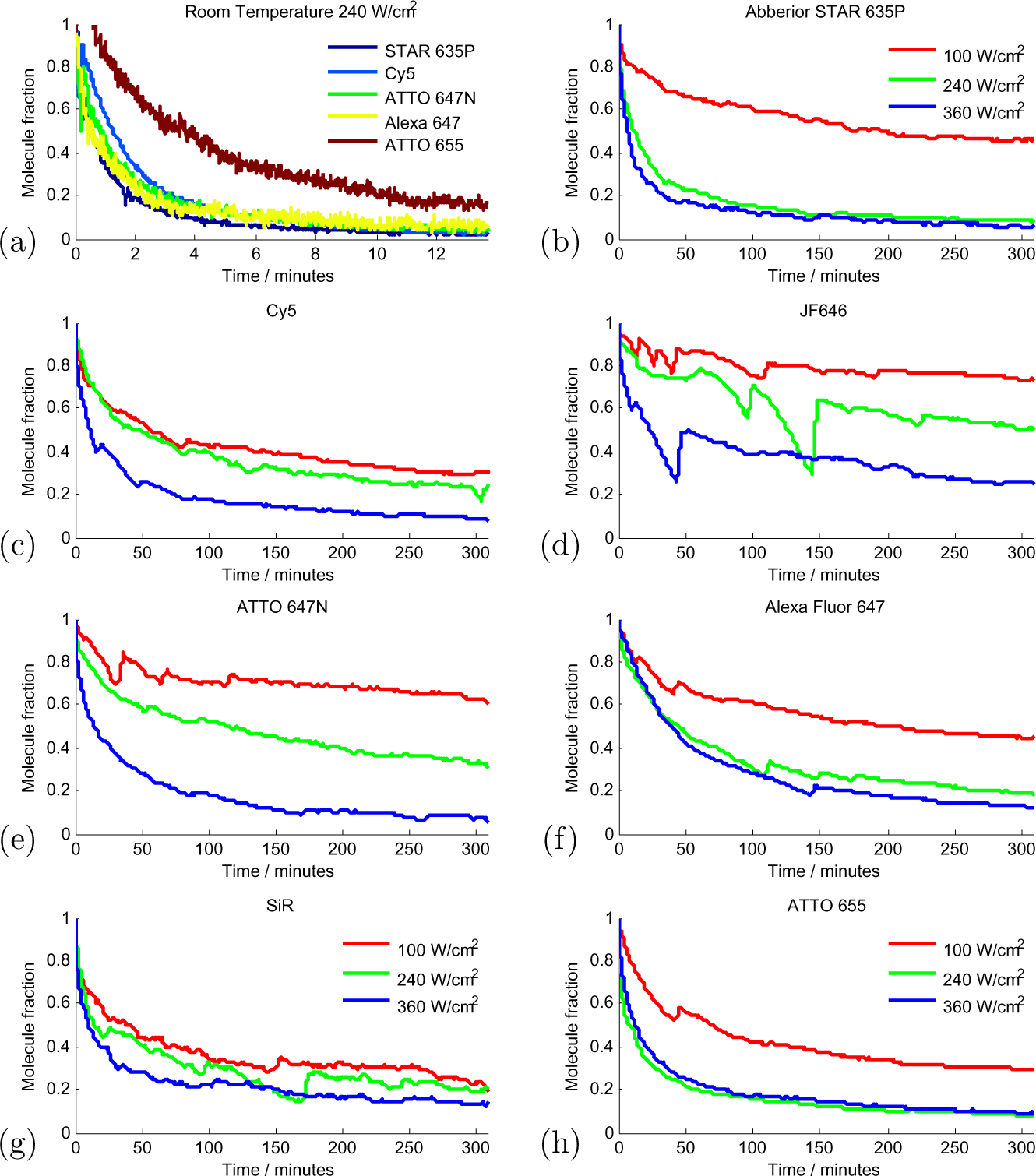
Photobleaching curves at room and cryogenic temperatures, as determined by the number of still fluorescing molecules per frame. **(a)**: Room temperature measurement of Abberior STAR635P, Cy5, ATTO 647N, Alexa Fluor 647 and ATTO 655 at 240 W/cm^2^ excitation intensity in the sample plane over 13 minutes. **(b)-(h)**: Decay curves at ~89 K and 3 different intensities 100 W/cm^2^ (red), 240 W/cm^2^ (green) and 360 W/cm^2^ (blue) for: **(b)** Abberior STAR635P, **(c)** Cy5, **(d)** JF 646, **(e)** ATTO 647N, **(f)** Alexa Fluor 647, **(g)** SiR and **(h)** ATTO 655. Curves are smoothed with a moving average filter with a span of 200 seconds. The large jumps in the curves are due to axial drift/defocusing, which required manual refocusing while imaging over several hours. Legends for **(b)-(h)** are identical.

In each image, the number of detected photons from a given molecule can be estimated, and when summing this number over all frames, the total photon yield per molecule till photobleaching is obtained (see Experimental Section for details). The photon yield distribution of ATTO 647N at room temperature at 240 W/cm^2^ excitation intensity shows an exponential distribution (Figure 2a). Thus, we fitted these photon yield histograms with an exponential decay function, *f* (*N*) = *A* · exp(−N/γ). Before fitting, we excluded the first bin because it can contain falsely identified weakly fluorescent signals. For ATTO 647N at room temperature, the mean photon yield from the exponential fit is γ = 2.1 ± 0.1 · 10^4^ photons, the uncertainty here is the 95% confidence interval of the fit. At cryogenic temperatures of ~ 89 K, the mean photon yield increases by a factor of 240 to γ = 5.1 ± 0.5 · 10^6^ photons for ATTO 647N at 240 W/cm^2^ excitation intensity. The mean photon yields for all the tested dyes at cryogenic temperatures range from γ = 2.9±0.8·10^6^ for SiR to γ = 6.5±0.3·10^6^ for JF 646 at 100 W/cm^2^ excitation intensity. This implies a mean theoretical localization precision of around 0.1 nm using equation 1 with a wavelength of λ = 685 nm and NA = 0.7. It is also possible to find the localization precision by calculating the standard deviation of the localizations over a few frames and scaling this by the amount of photons captured. For Abberior star 635P at 100 W/cm^2^ excitation intensity, we found an average standard deviation of 12 nm over 10 frames for 9 different molecules with an average total photon yield of 7 · 106 photons. Scaling with the ratio 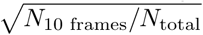 gives a mean localization precision of 0.3 nm which is on the same order of magnitude as the theoretical localization precision. This implies that all tested dyes are suitable for high resolution cryogenic super-resolution microscopy, as this is already smaller than the linker length used to attach fluorescent dyes to biological molecules of interest [15].

**Figure 2.**
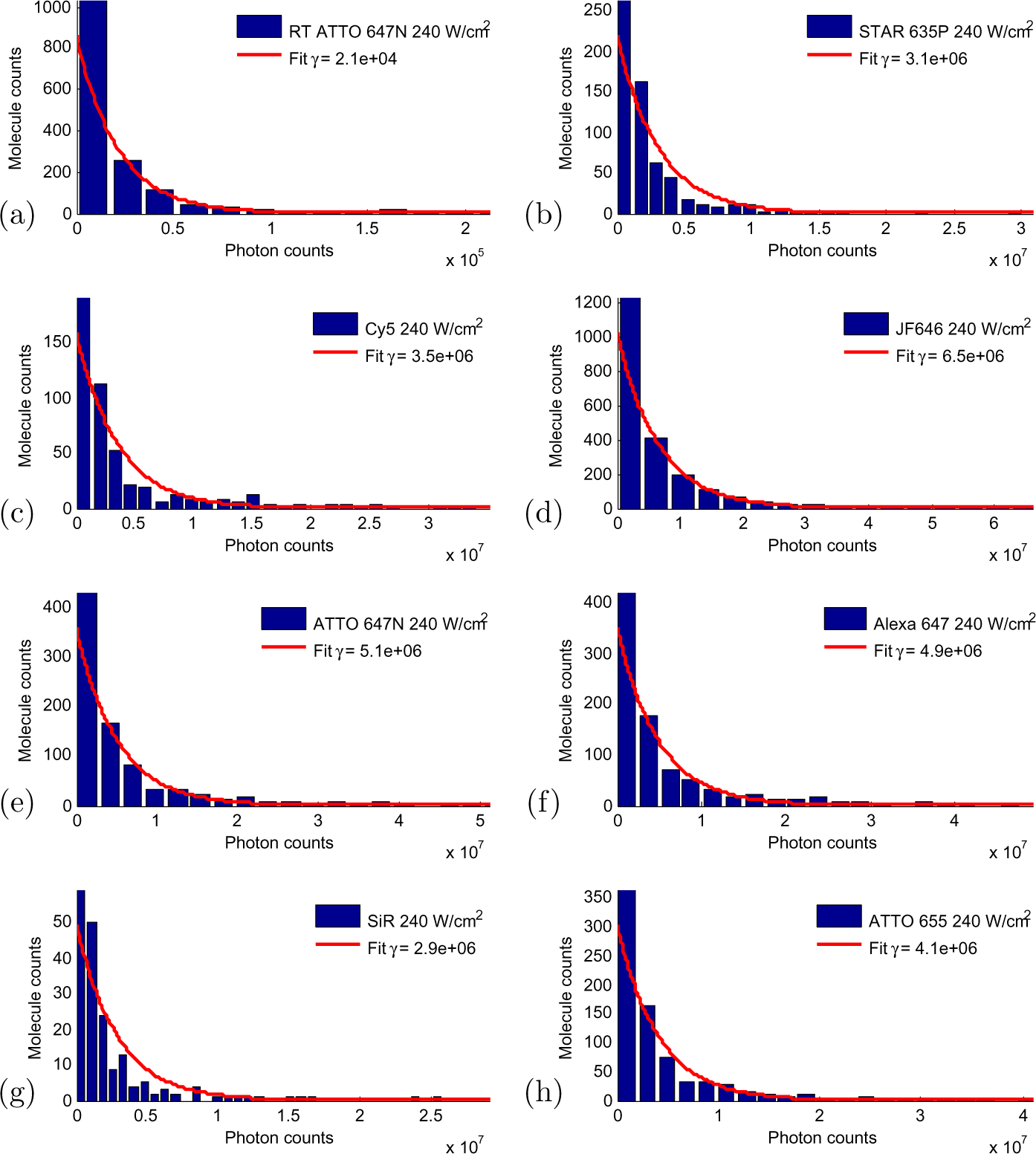
Histogram of photon counts fitted with an exponential decay. **(a)**: Room temperature measurement of ATTO 647N at 240 W/cm^2^ excitation intensity. Cryogenic measurements (~89 K) at 240 W/cm^2^ excitation intensity for **(b)** Abberior STAR635P, **(c)** Cy5, **(d)** JF 646, **(e)** ATTO 647N, **(f)** Alexa Fluor 647, **(g)** SiR and **(h)** ATTO 655.

To better compare the results between different dyes, we corrected the measured photon yield numbers by taking into account the different overlap between a dye’s emission spectrum and the transmission spectrum of the optical emission filters used in our experiment. Thus, we calculated the photon yield that one would observe if the emission filter would perfectly match the emission spectrum of a given dye. Again, we find that in most cases the photon yield γ is larger for lower excitation intensities (Figure 3a). The highest determined mean photon yield is γ = 11.1 ± 0.9 × 10^6^ photons for the dye Abberior STAR635P at 100 W/cm^2^ excitation intensity. The maximum absolute photon yield that we observed (over all detected molecules for a given dye) ranged between 7.9 · 107 photons for an SiR dye molecule and 3.6 · 108 photons for a JF 646 dye molecule.

**Figure 3.**
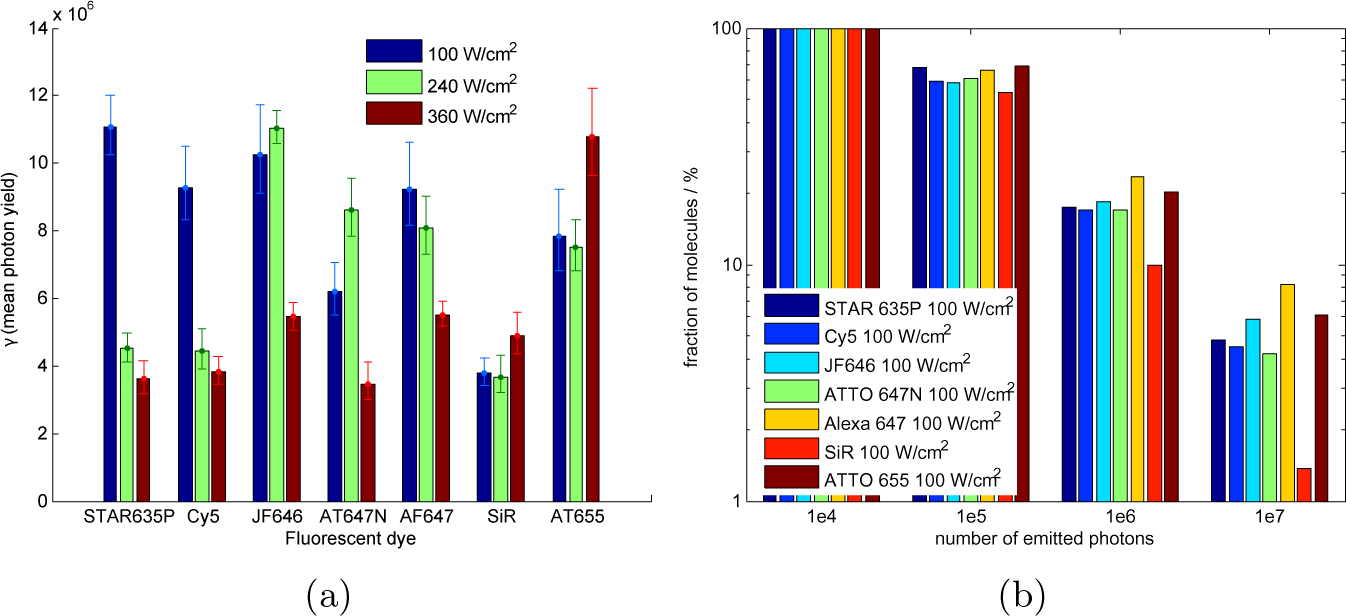
**(a)**: Mean photon yield γ and 95% confidence interval from the exponential fits of the photostability decay curves of γ dyes for 3 different intensities at cryogenic temperatures (~89 K). **(b)**: Cumulative histogram of the number of emitted photons per molecule at 100 W/cm^2^ excitation intensity and ~89 K. All molecule numbers were referenced against the number of molecules that survived the first minute of measurement time (100%, left columns in bar plot).

A cumulative histogram of emitted photons per molecule at 100 W/cm^2^ excitation intensity shows the distribution of detectable photons till photoble-aching at cryogenic conditions (Figure 3b). These histograms were normalized to the total molecule count (first left-hand columns in the figure) which is the sum of all molecules that were fluorescent for at least one minute after start of the measurement.

Finally, we can use equation 1 to calculate the minimal required photon yield per molecule to achieve a localisation precision of 1 nm, which is 6·10^4^ photons for our system. Typically, 80% of all molecules emit 6·10^4^ photons or more for the tested dyes and different excitation intensities (Figure 4a). The chance of finding a molecule with a localisation precision equal to or better than 0.1 nm (≥ 6 · 10^6^ photons) was found to be higher for a lower excitation intensity (100 W/cm^2^) for all dyes except ATTO 655 (Figure 4b). The dyes with the highest probability of achieving ≤0.1 nm localisation precision are Alexa Fluor 647 (11.1%), ATTO 655 (14.4%), and JF 646 (8.7%). The molecules emitting these amounts of photons were fluorescing during almost the entire experiment suggesting that the photon count from these molecules could be futher increased by extending the measurement time.

**Figure 4.**
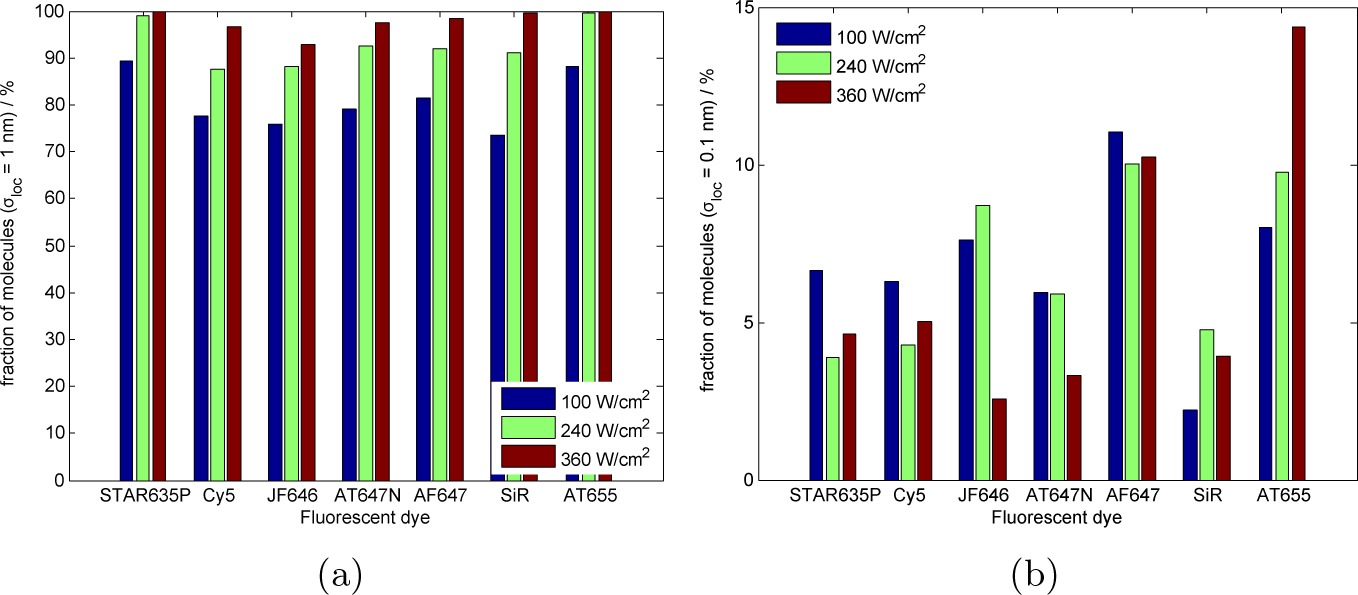
Fraction of molecules achieving a certain theoretical localisation precision for 7 different dyes and 3 excitation intensities at cryogenic temperatures (~89 K). **(a)**: ~1 nm (6 · 10^4^ photons) **(b)**: ~0.1 nm (6 · 10^6^ photons).

## 3 Conclusions

The mean photon yield of ATTO 647N (uncorrected for spectra) of 5 · 10^6^ photons at 240 W/cm^2^ and 3 · 10^6^ photons at 360 W/cm^2^ agrees well with the result found by Li et al. (3.5 · 10^6^ photons at 300 W/cm^2^) [7]. The maximal photon yield at liquid nitrogen temperatures is lower than at liquid helium temperatures, with only ~ 5% emitting 10^7^ photons or more compared to > 20% at liquid helium temperatures as found by Weisenburger et al. [16]. Although the photon yield is lower, liquid nitrogen is more suitable for fluorescence microscopy as phonons are frozen out at liquid helium temperatures leading to a significant narrowing of the excitation and emission lines [17], making efficient excitation of a fluorophore difficult.

All tested dyes show good performance for cryogenic localization, achieving an average theoretical localization precision around 0.1 nm from 3.5 · 10^6^ to 11 · 10^6^ photons. This is a more than hundred-fold increase in photon yield over room temperature, leading to a 10 fold increase in localization precision. From this study it seems that JF 646, Alexa Fluor 647, and ATTO 655 give the highest photon yields resulting in the best localization precision. It is assumed that due to the low density, most identified molecules are indeed single-molecules, which is also corroborated by the observed single-step bleaching of almost all identified molecules.

At ~89 K we found that the mean photon yield of the best dye was 3.1 times higher than the dye with the lowest mean photon yield. This means that there is a difference factor of 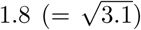 between the localization precision of the tested dyes. However, photon yield at cryogenic temperatures is not the only factor to be taken into consideration for dye selection. Other important factors are ease of biological labeling, solubility, cell permeability, net charge, extinction coefficient, stability, and non-specific staining.

This study has only addressed photon yield and its impact on localization of single molecules. To perform super-resolution measurements on biological samples, it is still necessary to induce sparsity so that individual molecules can be localized. One obvious way to do that could be to use PALM or dSTORM. All the dyes that were studied here do not show significant photoswitching which could be used for dSTORM. Also, we did not see any photoactivation of PA-JF 646 [18] at cryogenic conditions, although this dye shows excellent photoactivation at room temperature. Thus, besides for maximum photon yield, future work has also to screen for photoswitching ability at cryogenic temperatures. Besides organic dyes, promising candidates might be fluorescent proteins. For example, it is known that the fluorescent protein PA-GFP can be activated under cryogenic conditions [9], indicating that cryo-PALM may be feasible using this protein. Another method could be to induce sparsity by using polarization-dependent excitation and stimulated emission depletion [19, 20], although this will require a much more demanding experimental setup.

## 4 Experimental Section

### 4.1 Cryostat

The microscope used in this study is built solely for cryogenic Liquid Nitrogen (LN2) temperature measurements and has already been extensively described earlier [7]. Samples coated on 200 *μ*m thick quartz cover slips are mounted in the cryostat and are imaged through a 0.5 mm thick quartz window to maintain vacuum. Quartz is used instead of glass due to its reduced autofluorescence. The sample chamber is pumped to a pressure ≤ 1.6 · 10^−5^ mbar. The cryostat is filled with LN2 and the temperature is left to stabilize for one hour. Once stabilized, it is possible to measure for more than 5 hours with a temperature between 89-93 K and a typical temperature change of 0.1 K/hour.

**Figure 5.**
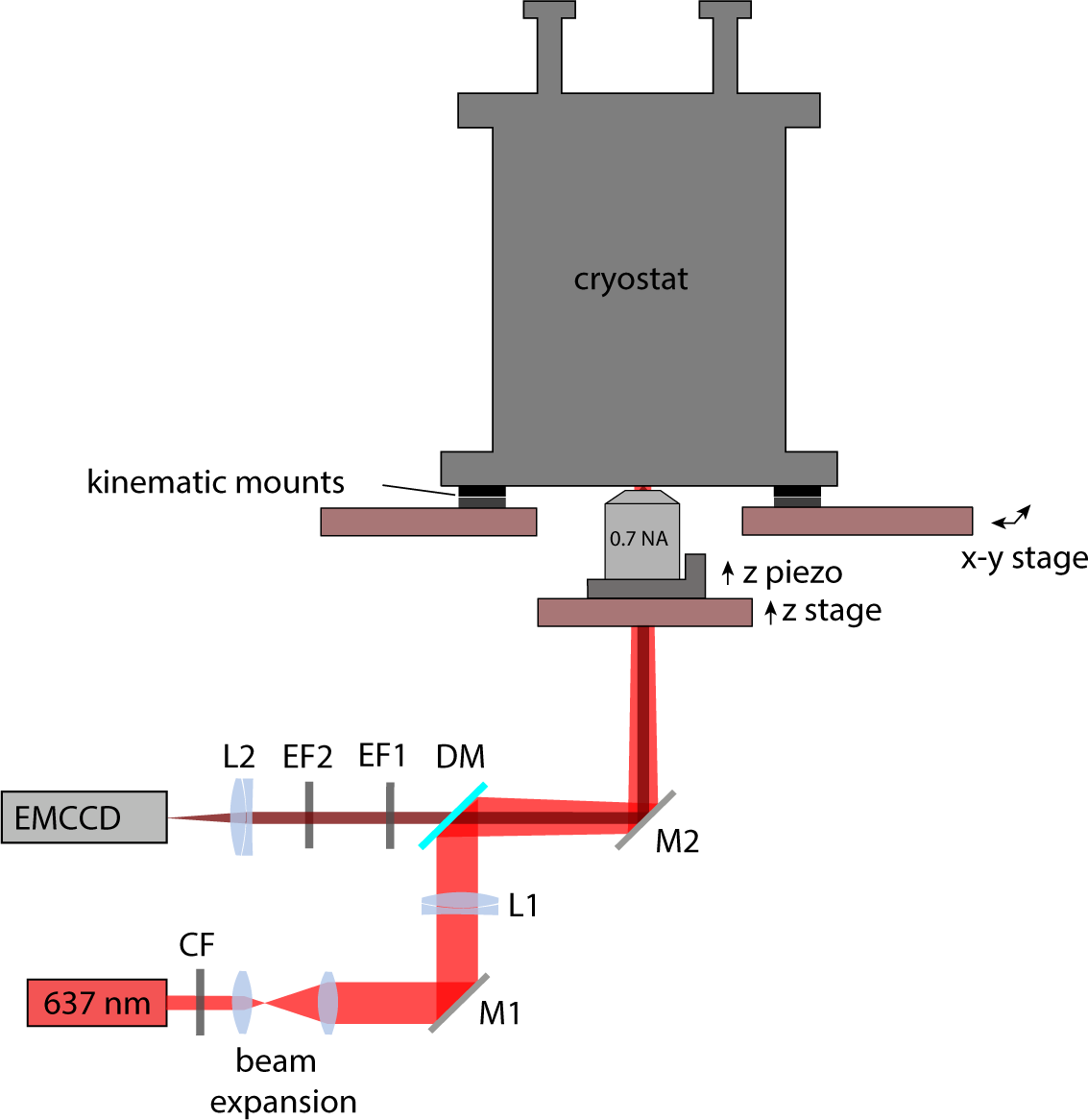
Cryo-fluorescence microscope with the sample mounted in the vacuum environment of the cryostat. Imaging is done with a long working distance 0.7NA objective onto an EMCCD camera. The excitation laser is focused onto the back focal plane of the objective and illuminates the field of view of the camera.

### 4.2 Fluorescence microscope

The whole cryostat is mounted on a motorized x-y stage, and the objective (LUCPLFLN 60x/0.7NA, Olympus) is mounted on both a motorized z stage and a piezo z-stage (PIFOC, PI). The excitation and emission path is that of a typical epi-fluorescence microscope (Figure 5). The red 637 nm continuous wave excitation laser (OEM-SD-637-500, CNI) is filtered by a clean-up filter CF (HC 640/14, Semrock), expanded by a telescope (Plano convex f=30 mm, f=100 mm, Thorlabs), and focused onto the back focal plane of the objective by lens L1. The dichroic mirror DM (Di01-R405/488/532/635, Semrock) reflects the excitation laser and transmits the fluorescence from the sample. The fluorescence is filtered by two emission filters EF1 (FF01-446/510/581/703, Semrock) and EF2 (ET 685/50, Chroma) and focused by lens L2 (AC508-180-A-ML, Thorlabs) onto an EMCCD camera (Ixon Ultra 897, Andor). Excitation intensities are estimated by measuring the power of the parallel beam after the objective with a power meter and dividing it by the illuminated area in the sample plane.

We investigated the dyes; Abberior STAR635P, Cy5, JF 646 (spectrum from [21]), ATTO 647N, Alexa Fluor 647, SiR and ATTO 655, which have very similar excitation and emission spectra (figure 6). The emission spectra are scaled to their values at 637 nm excitation, and they are plotted along with the transmission curve of the emission filter combination used in our experiment (figure 6b). To correct for the effect of the emission filter and imperfect excitation wavelength, we up-scaled photon count numbers by the scaling factor

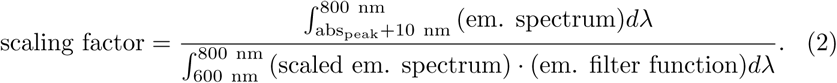

In the denominator, we start the integration only 10 nm after the absorption peak to model the gradual transition between reflection and transmission of a dichroic mirror. In order to obtain the compensated photon counts, the raw photon count number is multiplied by this scaling factor for each dye. The scaling factor ranges from 1.7 for JF 646 to 2.5 for Abberior STAR635P.

**Figure 6.**
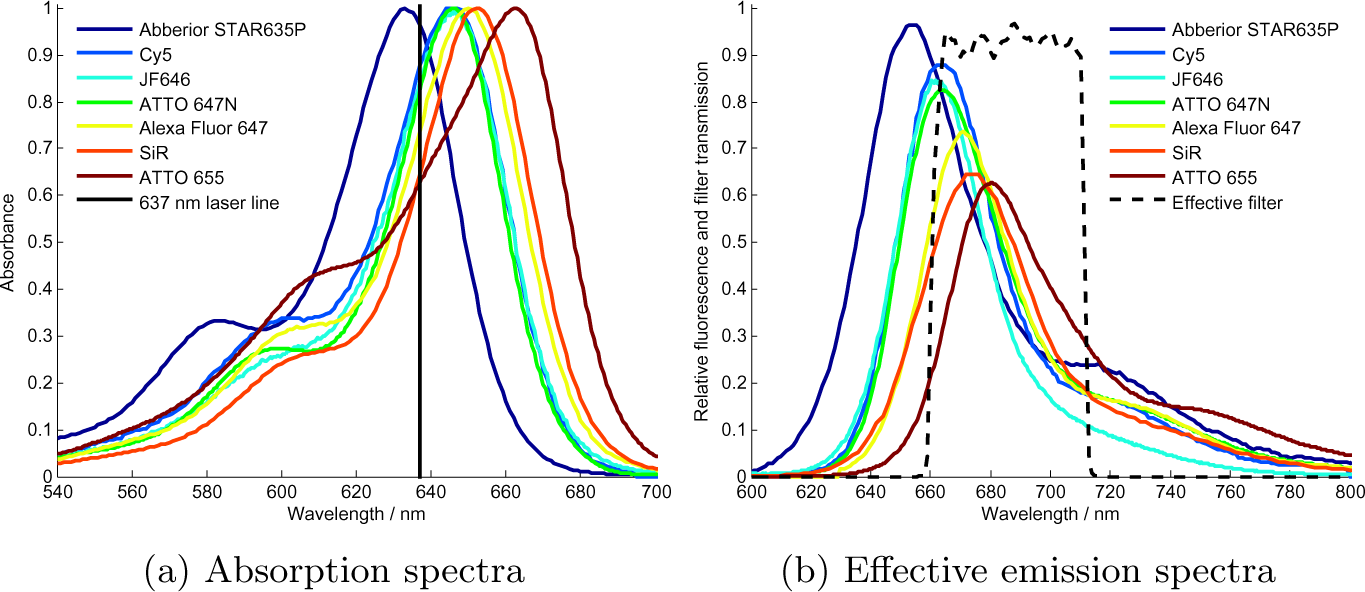
Absorption and emission spectra of Abberior STAR635P, Cy5, JF 646, ATTO 647N, Alexa Fluor 647, SiR and ATTO 655. The emission spectra are normalized with excitation at 637 nm. The effective filter is the combined action of the dichroic mirror, quad-band filter and ET685/50 bandpass filter.

### 4.3 Sample preparation

Initially, each organic dye is diluted with distilled, de-ionized and filtered water to a 100 pM concentration. 5 *μ*L of this dilution is deposited on a quartz slide and spin-coated at 3000 RPM for 45 seconds. The quartz slide is imaged at room temperature to determine the density of molecules in the 136.5 *μ*m × 136.5 *μ*m field of view. We aimed at having ~400 single molecules in the field of view to balance sparsity whilst retaining enough molecules after extensive bleaching. Dye concentrations were adjusted between 25 pM and 500 pM to achieve the desired number of initial molecules. At cryogenic temperatures, the initial amount of single molecules was found to be higher, 400-1200 instead, because at room temperature many molecules would have already been bleached during focusing, thus lowering the amount of molecules. Concentrations were not compensated for this effect but kept at 200-400 localizations at room temperature to maintain consistency between measurements.

### 4.4 Image acquisition

For each cryogenic photon yield measurement, the full time duration over which the cryostat is stable is used, to maximize the total amount of captured photons. In total, 18,000 frames with 1 second exposure time are recorded, resulting in 130-220 photons per molecule per frame. Each frame has a readout time of 25 ms, resulting in a total measurement of ~5 hours. All data is acquired with an EM gain of 100, and an EMCCD sensor temperature of −45 °C. Although the system is very stable, there is still a typical axial drift of 3 *μ*m over 5 hours causing defocusing. To compensate for this, it is necessary to manually refocus the system periodically. We found that refocusing every 10 minutes during the first hour and then after every hour thereafter is sufficient.

### 4.5 Image analysis

The Analog-Digital Units (ADU) of the camera images can be converted into photons when knowing the bias *b*, EM gain *G*, and specific sensitivity *s* of the camera, by using the equation

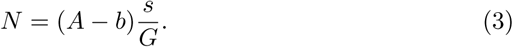

The amplitude of the Gaussian fit to a single molecule image was used to calculate the total number of emitted photons from a molecule in each frame. Recently, it has been shown that 2D Gaussian fitting can lead to a 10-30% under-estimation of the photon counts [22], but we did not take this into account here. After tracking each molecule over the course of the whole measurement, the total number of emitted photons per molecule was determined.

Molecule tracking was done using TrackNTrace, an open-source MATLAB framework for localizing and tracking molecules [23]. TrackNTrace identifies molecules in two steps. We used a cross-correlation for the candidate detection step, and a 2D Gaussian fitting for the refinement step. The localized molecules were tracked using a nearest neighbor approach.

The tracking procedure was applied twice, where the initial tracking is done with no frame gaps, meaning that blinking leads to new tracks. This results in reliable data to calculate the average drift from frame to frame. The data is then adjusted for this average drift and tracked again, but now frame gaps are allowed (up to the total measurement duration). This allows molecules to blink but still be identified as one and the same molecule. This method reduces the chance of identifying distinct molecules as a single molecule. Combined with the low density of molecules, we assume that almost all identified molecules are single.

## 5 Acknowledgments

We thank Luke Lavis (HHMI, Janelia Farm) for providing us with the PA-JF 646 and JF 646 dyes. B.R. and C.N.H. acknowledge European Research Council grant no. 648580. W.L., I.G. and J.E. acknowledge financial support by the Deutsche Forschungsgemeinschaft (DFG, project A06 of the SFB 860).

